# Dynamic representation of taste-related decisions in the gustatory insular cortex of mice

**DOI:** 10.1101/2019.12.16.878553

**Authors:** Roberto Vincis, Ke Chen, Lindsey Czarnecki, John Chen, Alfredo Fontanini

**Affiliations:** Department of Neurobiology and Behavior, Stony Brook University, Stony Brook, NY, 11794, USA; Graduate program in Neuroscience, Stony Brook University, Stony Brook, NY, 11794, USA; Department of Biological Science and Program in Neuroscience, Florida State University, Tallahassee, FL, 32306, USA

**Keywords:** gustatory cortex, decision making, temporal dynamics, licking

## Abstract

Research over the past decade has established the gustatory insular cortex (GC) as a model for studying how primary sensory cortices integrate multiple sensory, affective and cognitive signals. This integration occurs through time varying patterns of neural activity. Selective silencing of GC activity during specific temporal windows provided evidence for GC’s role in mediating taste palatability and expectation. Recent results also suggest that this area may play a role in decision making. However, existing data are limited to GC involvement in controlling the timing of stereotyped, orofacial reactions to aversive tastants during consumption. Here we present electrophysiological, chemogenetic and optogenetic results demonstrating the key role of GC in the execution of a taste-guided, reward-directed decision making task. Mice were trained in a taste-based, two-alternative choice task, in which they had to associate tastants sampled from a central spout with different actions (i.e., licking either a left or a right spout). Stimulus sampling and action were separated by a delay period. Electrophysiological recordings of single units revealed chemosensory processing during the sampling period and the emergence of task-related, cognitive signals during the delay period. Chemogenetic silencing of GC impaired task performance. Optogenetic silencing of GC allowed us to tease apart the contribution of activity during the sampling and the delay periods. While silencing during the sampling period had no effect, silencing during the delay period significantly impacted behavioral performance, demonstrating the importance of the cognitive signals processed by GC during this temporal window in driving decision making.

Altogether, our data highlight a novel role of GC in controlling taste-guided, reward-directed choices and actions.

## INTRODUCTION

The gustatory cortex (GC), a subregion of the insular cortex, has traditionally been investigated for its function in processing taste identity [1]. In the past decade, studies in alert animals significantly changed the classic view, establishing a role for GC in dynamically representing affective, multisensory and cognitive signals associated with the experience of eating [2–4]. Time varying patterns of firing activity in GC are important for the perception and learning of taste value [5–7], for multisensory integration in the context of flavor and taste expectation [8–12], and for guiding food-directed behaviors on the basis of food-predictive cues [13–15].

Recent experiments indicated that GC may also be involved in mediating decisions based on gustatory cues. Electrophysiological recordings and optogenetic manipulations in rats consuming tastants demonstrated that GC activity is instructive of ingestive decisions [16]. Indeed, sudden changes in ensemble activity occurring during the time course of a response correlated with and determined the onset of gapes – aversive reactions aimed at expelling highly unpalatable tastants [16]. The function of GC is not limited to naturalistic consummatory decisions involving stereotyped, orofacial reactions to aversive tastants. Single unit recordings in an operant task classically used to study perceptual decision making (i.e., a taste-based, two-alternative choice task [2-AC]) suggested that neurons in GC may encode taste-guided, reward-directed choices and actions [17]. However, the extent to which activity in GC contributes to driving reward-directed choices in a 2-AC task is currently unknown. Furthermore, it is not established whether GC contributes to decision making by exclusively representing chemosensory information (i.e. sensory evidence necessary for decisions), or by encoding also cognitive variables such as planning for specific behavioral choices and actions.

In this study, we addressed these unresolved issues by recording and manipulating GC activity in the context of a taste-based, two-alternative choice task optimized for the investigation of sensory and task-related variables. We designed a 2-AC task in which pairs of gustatory stimuli of opposite categories (sweets and bitters) sampled from a central spout were rewarded with water delivered at two lateral spouts. The task featured a delay period, specifically introduced to better resolve activity anticipating decisions and actions [18]. We recorded GC neurons’ spiking activity in well trained, head-restrained mice. Analysis of single unit and population activity revealed a progression from chemosensory coding to the representation of task-related variables. Specifically, we observed that GC neurons encode information about the action-predictive value of tastants, and about planning of an imminent behavioral choice during the delay period. The behavioral significance of this task-related activity was validated with optogenetic silencing of GC, which demonstrated that interfering with activity during the delay epoch, but not taste sampling, significantly reduced behavioral performance.

Our results show that GC neurons dynamically encode multiple variables associated with a perceptual decision making task, and demonstrate that activity during the period preceding a taste-guided, reward-directed choice is instructive of behavior. This evidence significantly changes our understanding of the function of GC in taste, demonstrating its role as a key node for gustatory decision making.

## RESULTS

### Performance in a taste-based, two-alternative choice task

We trained head-restrained mice to perform a taste-based, two-alternative choice (2-AC; **Figure 1**) task in which sensation (i.e. sampling of taste stimuli) and action (i.e. lick left or right spout for reward) are separated by a delay epoch (~ 2 s) (**Figure 1B**). Mice were trained to sample 2 μl of one out of four taste stimuli (sucrose [100 mM], quinine [0.5 mM], maltose [300 mM] and sucrose octaacetate [0.5 mM]) delivered from a central spout at each trial, and to associate pairs of tastants with different actions (**Figure 1A-B**). After a delay epoch, initiated by the retraction of the center spout, two lateral spouts advanced and mice could lick towards the left or right lateral spout to receive a small drop of water reward (3 μl). Mice were trained to associate sucrose (S) and quinine (Q) with reward from the left spout, and maltose (M) and sucrose octaacetate (SO) with reward from the right spout. In this configuration, each action (left or right lick) was paired with two tastants with opposite hedonic value and different taste quality, rendering mice unable to solve the task by simply generalizing for taste palatability or quality.

**Figure 1.**
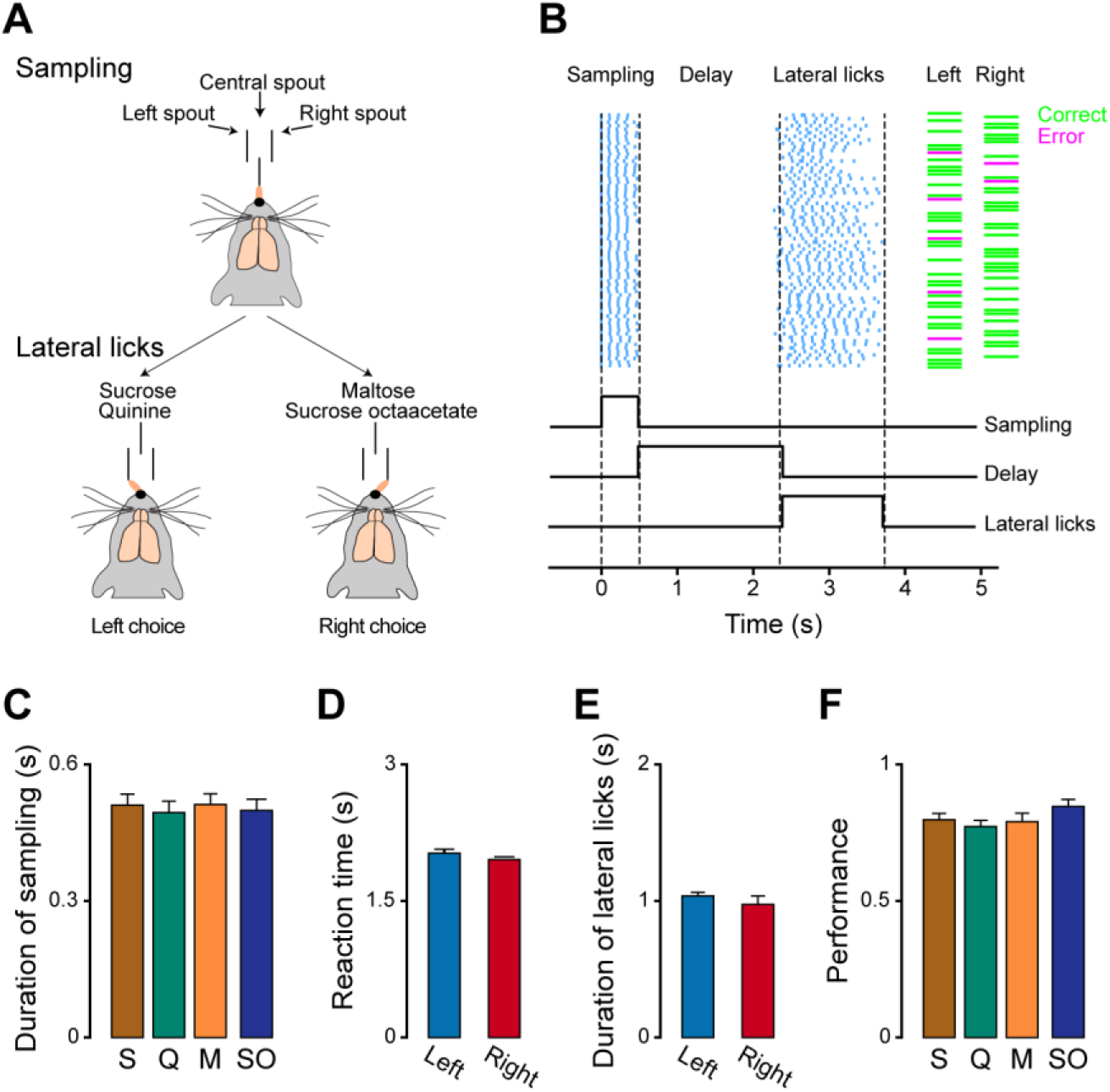
Taste-based, two-alternative choice task. **A**, Diagram showing a head-fixed mouse sampling tastants from a central spout and responding with appropriate licking. **B**, Top panel: representative raster plots of licking activity during a behavioral session. Each row represents a single trial, and each cyan tick represents a lick. The green horizontal bars represent correct trials and the magenta horizontal bars represent errors. Bottom panel: schematic diagram of the taste-based, 2-AC with its three epochs: sampling, delay and lateral licks. **C**, Bar plots showing the average duration of taste sampling (i.e. how long mice licked to the central spout during the sampling epoch) for each stimulus (sucrose: S, quinine: Q, maltose: M, sucrose octaacetate, SO). **D**, Bar plots showing the average reaction time from the end of taste sampling to the first lateral lick for left (blue) and right (red) trials. **E**, Bar plots showing the duration of lateral licks for left (blue) and right (red) correct trials. **F**, Bar plots showing the average of behavioral performance (fraction of correct choices) for the four gustatory stimuli. In **C-F** bar plots (n = 16 mice), error bars represent standard error of the mean (SEM).

Upon learning the task, mice showed no bias in the performance. The average duration of the sampling (i.e. the time during which a mouse licked the central spout to sample the tastant) was 0.50 ± 0.02 s, the average licking frequency was 8.65 ± 0.16 Hz. No significant difference in sampling duration or licking frequency was observed for the four tastants (n = 16, one-way ANOVA, for sampling duration, F(3,60) = 0.12, p = 0.94, **Figure 1C**; for licking frequency, F(3,60) = 0.04, p = 0.99). The reaction time for left trials (measured as the interval between the last lick for the center spout and the first lick for a lateral spout) was comparable to that for right trials (n = 16, 2.02 ± 0.04 s vs 1.95 ± 0.03s, student’s t-test, t_(30)_ =1.25, p = 0.21, **Figure 1D**), and mice showed similar licking duration and frequency to each lateral spouts (n = 16, left vs right, duration:1.04 ± 0.03 s vs 0.97 ± 0.06 s, student’s t-test, t_(30)_ = 0.90, p = 0.37, **Figure 1E;** frequency: 7.22 ± 0.15 vs 7.44 ± 0.38 Hz, student’s t-test, t_(30)_ = −1.06, p = 0.30), indicating lack of any lateral bias. Finally, mice showed similar behavioral performance for each of the four tastants (n = 16, one-way ANOVA, F(3,60) = 1.5, p = 0.22, **Figure 1F**), denoting that they could learn the contingency for each tastant, and further confirming the absence of any bias toward one or more specific tastants used in the task.

### Taste classification during sampling and delay epochs

Single unit spiking activity was recorded with movable bundles of 8 tetrodes unilaterally implanted in GC of mice performing the 2-AC task at criterion (**Supplementary Figure 1A**). Neural activity, licking activity as well as orofacial movements were simultaneously recorded. Given the involvement of GC in representing taste [19, 20], we first analyzed activity evoked by S, Q, M, and SO during the sampling epoch. Spiking activity was aligned to the first lick at the central spout (time 0, detection of the taste, **Figure 2A**), and analyzed for a 500 ms temporal window (sampling epoch; **Figure 2A**). As expected, a sizable portion of GC neurons changed their firing rate following the licking of a gustatory stimulus and had significantly different responses to the four tastants (**Figure 2B**). Specifically, we observed that 33.6% (72/214) of recorded neurons were modulated by at least one of the four tastants (**Figure 2C**). Of these taste responsive neurons, 73.6% (53/72) were modulated by S, 63.8% (46/72) by Q, 84.7% (61/72) by M and 66.7% (48/72) by SO (**Figure 2D**).

**Figure 2.**
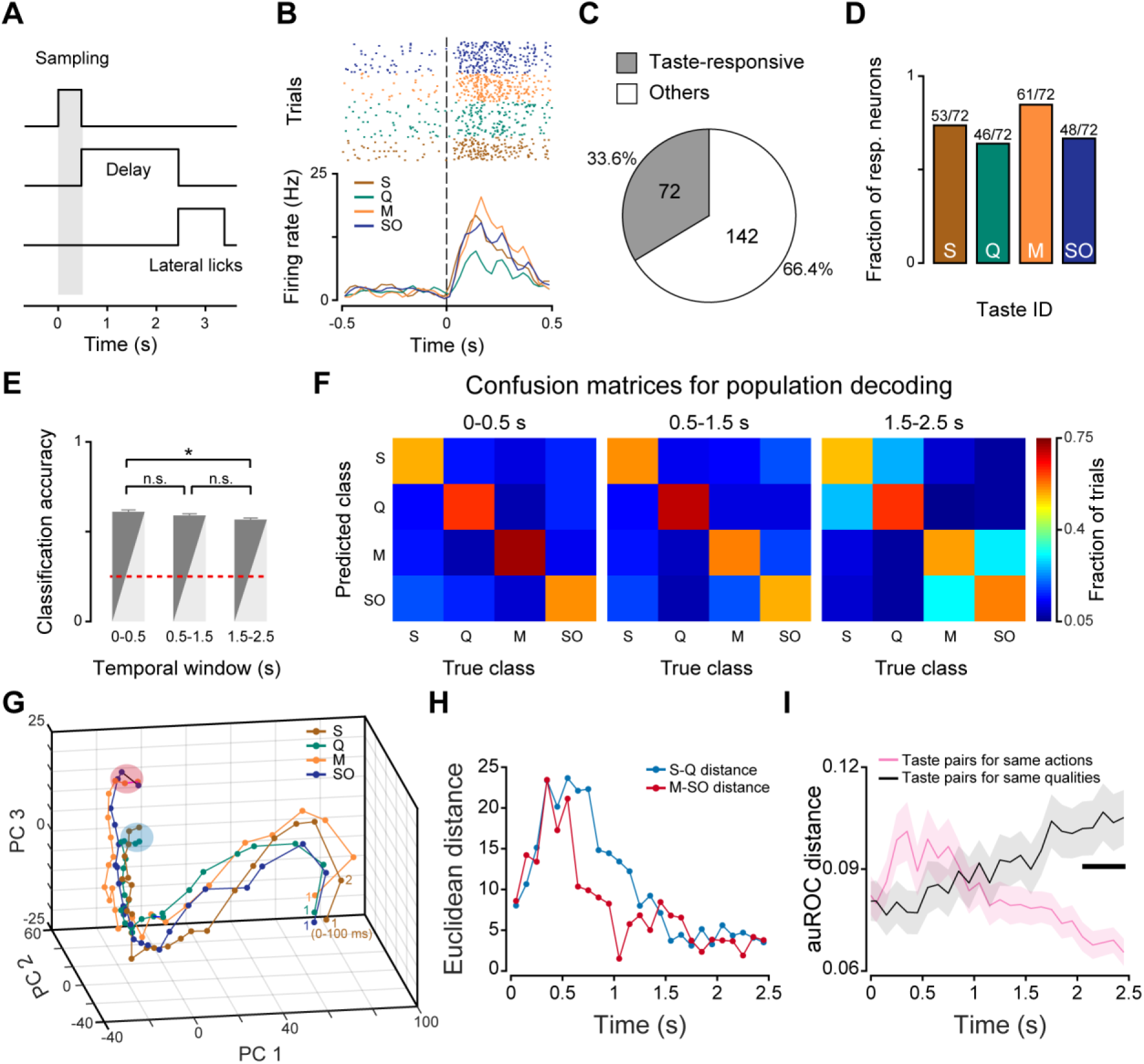
Taste representation in GC. **A**, Schematic showing the trial structure. The gray bar represents the temporal window (500 ms, sampling epoch) in which we analyzed taste responses. Time 0 represents the first lick to the central spout. **B**, Raster plot and PSTH for a representative neuron showing responses to the four taste stimuli. Dashed lines at time 0 represent the first lick to the central spout. **C**, Pie chart showing the proportion of taste responsive (gray) and non-responsive (white) neurons. **D**, Bar plots showing the fraction of taste responsive neurons modulated by each of the four gustatory stimuli used. **E**, Bar plots showing population decoding accuracy for three different temporal windows. Time 0 is the first lick to the central spout. Temporal window from 0 to 0.5 s: sampling epoch; windows from 0.5 to 1.5 s and 1.5 to 2.5 s: delay epoch. Bars represent the mean, and error bars represent SEM. One-way ANOVA and post hoc Tukey’s HSD test, * p<0.5; n.s. indicates not significant. **F**, Confusion matrix showing decoding performance for each tastant in the three different temporal windows (left, 0-0.5 s; middle, 0.5-1.5 s; right, 1.5-2.5 s). **G**, Trajectories of population activity in PC space for responses to each of the 4 gustatory stimuli. “1” represents the first bin (i.e. 0-100 ms) following the first lick to the central spout. The blue and red shaded areas highlight the convergence at the end of the delay (2.2-2.5 s) of S/Q-evoked activity and M/SO-evoked activity respectively. **H**, Temporal profiles of Euclidean distance in PC space. Blue curve: Euclidean distance between S and Q-evoked trajectories; red curve: Euclidean distance between M and SO-evoked trajectories. **I**, Time course of pairwise difference in firing responses for different tastants. The magenta trace shows the average distance for pairs of tastants associated with the same actions. The black trace shows the average distance for pairs of tastants associated with same qualities. Shading represent SEM. The thick horizontal black bar represents times at which the auROC distance is significantly different across the two groups (t-test with p < 0.05/25).

Gustatory processing in GC is dynamic, and evidence from the literature suggests that responses may persist or emerge beyond the initial 500 ms sampling epoch [5]. To begin assessing the temporal dynamics of gustatory processing, we performed a population decoding analysis across sampling and delay epochs. We found that taste decoding was more accurate in the sampling epoch (0-0.5 s) compared to the later part of the delay epoch (1.5-2.5 s), indicating that taste decoding accuracy slightly decays during the delay (n = 181, see methods, decoding accuracy: 0.61 ± 0.01 in sampling epoch, 0.59 ± 0.01 and 0.57 ± 0.01 in the delay epoch; one-way ANOVA, F(2, 27) = 4.8, p = 0.016; post hoc Tukey’s HSD test, p <0.05, **Figure 2E**). In addition, we constructed confusion matrices for population decoding and characterized the classification performance for each taste. We found that compared to the sampling epoch or the first part of the delay (0.5-1.5 s), the decoder made more mistakes between tastants associated with the same actions (i.e. S and Q trials or M and SO trials) in the later part of the delay (1.5-2.5 s; **Figure 2F**). This observation suggests that neural activity evoked by tastants associated with the same action converges during the late phase of the delay epoch. To visualize temporal dynamics of population activity, we applied a principal component analysis (PCA, **Figure 2G**). Visual inspection of the trajectories of taste-evoked temporal dynamics reveals that S- and Q-evoked activity converged to the same small region in the PC space at the end of the delay (blue spot, **Figure 2G**), and that M- and SO-evoked activity converged to a distinct spot in the PC space (red spot, **Figure 2G**). The Euclidean distance in PC space between S- and Q-evoked activity or between M- and SO-evoked activity gradually decreased in the delay epoch (0.5 -2.5 s, **Figure 2H**). To confirm that activity becomes more similar for pairs of tastants associated with the same actions, we computed the pairwise distance in normalized firing rates evoked by each taste for each neuron (n = 214, see method). The distance for firing activity evoked by pairs of tastants associated with the same actions gradually decreased – reflecting an increase in the similarity of the responses. In contrast, the distance for pairs of tastants associated with the same taste quality (sweet vs bitter) gradually increased (**Figure 2I**).

Altogether, these data demonstrate that in the context of a perceptual decision making task, taste processing is not restricted to the sampling epoch, but continues throughout the delay period, and that GC categorizes tastants according to different criteria in different epochs. As time progresses, GC shifts from coding the chemosensory identity of tastants to firing more similarly for stimuli anticipating the same action.

### Action-related activity in the delay epoch

To further investigate neural activity during the delay epoch and identify neurons responsible for the changes seen in confusion matrices and pairwise distances, we computed each neuron’s preference for firing in anticipation of correct left or correct right licking using a receiver operating characteristic (ROC) analysis (see methods, [21]). A large group of neurons (41.6% of all recorded neurons, 89/214) showed a significant direction preference in their firing (**Figure 3B)**, with 57.3% (51/89) and 42.7% (38/89) of neurons having preference for the anticipation of leftward and rightward licking, respectively (**Figure 3C-D**). **Figure 3B** shows raster plots and PSTHs for two representative neurons, one with higher firing rate during the delay epoch in left trials (Neuron #1, leftward preference), and the other showing higher firing rate for right trials (Neuron #2, rightward preference). Direction selective firing could begin at any time during the delay period - i.e., from 2 seconds prior, to the moment of the lateral lick - as shown in the color coded population PSTH in **Figure 3D**. Inspection of the average direction preference for left and right trials (white traces superimposed to the color plot in **Figure 3D**) revealed that direction selectivity peaks right before the animal licks the lateral spouts.

**Figure 3.**
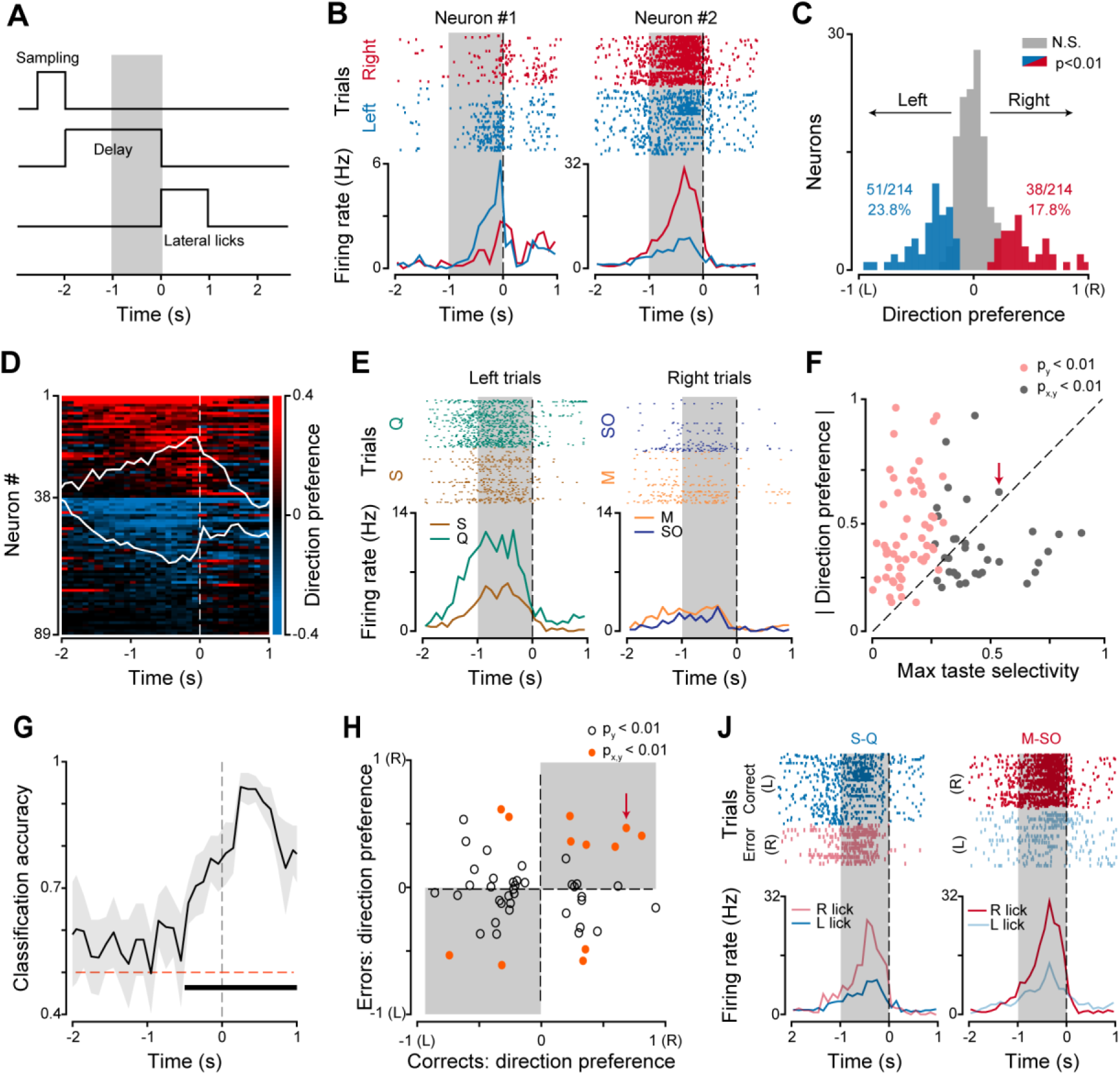
Preparatory activity in GC. **A**, Schematic of trial structure. The gray bar highlights the temporal window (1 s) used to analyze preparatory activity. Time 0 represents the first lick to the lateral spout. **B**, Raster plots and PSTHs of two representative neurons showing direction-selective, preparatory activity. The neuron on the left (Neuron #1) displays higher firing rates during the delay period preceding left licks (blue ticks and blue line for raster plot and PSTH, respectively); the neuron on the right (Neuron #2) displays higher firing rates in anticipation of right licks (red ticks and red line for raster plot and PSTH, respectively). Time 0 represents the first lick to the lateral spout. **C**, Histogram of direction preference during the delay epoch. Blue and red bars represent neurons with statistically significant direction preference for left- and right-correct trials, respectively. Gray bars represent neurons with no significant direction preference (similar firing rate between left and right correct trials). **D**, Heatmap showing the time course of direction preference. Each row represents a single neuron (only neurons with direction preference are shown). Time 0 is the first lick to the lateral spout. White traces superimposed on the heatmap represent the average direction preference for neurons with leftward (preference < 0, bottom) and rightward (preference >0, up) preference. **E**, Raster plots and PSTHs for one neuron showing preparatory activity and taste selectivity during the delay epoch. On the left (left trials), raster plot and PSTH for sucrose (S, brown) and quinine (Q, green) trials; on the right (right trials), raster plot and PSTH for maltose (M, gold) and sucrose octaacetate (SO, blue) trials. Time 0 is the first lick to the lateral spout. **F**, Scatter plot showing the relationship between max taste selectivity and the absolute value of direction preference. Each dot (pink and gray) represents a neuron with significant direction preference (p_y_ < 0.01); Gray dots represent neurons that also show taste selectivity during the delay epoch (p_x,y_ < 0.01). The gray dot with the red arrow represents the neuron shown in panel **E**. **G**, Time course of classification accuracy for correct and error trials. Time 0 represents the first lick to the lateral spout. The red horizontal dashed line represents classification accuracy at chance level (0.5). The thick horizontal black bar represents times with classification accuracy that is significantly higher than chance level (permutation test, p < 0.001). Shading represents the 99.5% confidence interval. **H**, Scatter plot showing direction preference in correct and error trials. Each dot represents a neuron with significant direction preference in correct trials. Orange points represent neurons that also show significant direction preference in error trials. Grey shaded areas highlight the quadrants in which neurons have comparable direction preference in correct and error trials regardless of the gustatory cue. The red arrow indicates the neuron shown in panel **J**. **J**, Raster plots and PSTHs for one neuron showing comparable direction preference in correct and error trials. Time 0 is the first lick to the lateral spout. Left, raster plots and PSTHs for correct (left licks, dark blue) and error (right lick, light red) trials in response to S and Q. Right, activity for correct (right lick, dark red) and error (left licks, light blue) trials in response to M and SO.

To determine whether these direction-selective neurons carried information regarding the chemosensory identity of specific tastants, we compared firing rates for S *vs* Q trials (left trials) or for M *vs* SO trials (right trials; **Figure 3E**). We found that 38.2% (34/89) of the neurons with significant direction preference also showed significant taste selectivity during the delay epoch (**Figure 3F**; gray dots). Plot of the maximum value for taste selectivity against the absolute value of direction preference revealed that the activity of the majority of neurons, 74.1% (66/89), was more strongly modulated by the anticipated direction of licking than by the chemosensory identity of the tastant (**Figure 3F**).

In principle, direction selective activity could be evoked either by the tastants (and reflect a taste recategorization according to each stimulus’ predictive value), by internal signals pertaining to the preparation/planning of a specific action or by a combination of both. To investigate these possibilities, we analyzed responses for correct and error trials for the same pairs of cues (e.g., correct: S and Q →left lick; error: S and Q → right lick). If GC was involved exclusively in taste recategorization, activity would depend just on gustatory cues, hence failing to differentiate error and correct trials. On the contrary, delay activity related to action planning would allow for the classification of correct and error trials for the same gustatory cues. A decoding analysis (**Figure 3G**) revealed that the delay activity in the population of neurons with direction selectivity can indeed differentiate between correct and error trials. Classification of correct and errors peaked short after the action (peak accuracy = 0.94, 0.25 s after lateral licking), but was already significant in the delay period (−0.5 to 0 s, permutation test with p <0.001). This classification performance was related to neurons with comparable direction preference regardless of the gustatory cue (grey shading in **Figure 3H**), like the one shown in **Figure 3J.** Not all direction selective neurons behaved like the one in **Figure 3J**; some neurons represented pairwise similarities between S and Q (or M and SO) regardless of action (unshaded area in **Figure 3H**, and **Supplementary Figure 2A**) indicating that GC can also represent taste recategorization and hence adopt a mixed coding scheme.

Direction preference and preparatory activity in the delay epoch may be related to specific orofacial movements anticipating left and right licking. To address this, we analyzed videos of the orofacial region during the delay period. Visual inspection of traces extracted from the video analysis (**Supplementary Figure 3B-C**) suggest that preparatory movements during the delay epoch were similar for left and right trials. ROC analysis confirmed that orofacial activity in left and right trials during the delay period was comparable for all the sessions analyzed (**Supplementary Figure 3D**). Thus, it is unlikely that the neural activity during the delay epoch relates to differences in orofacial movements.

Altogether, the results reveal that during the delay epoch a large fraction of GC neurons can show firing rate modulations in anticipation of a specific licking direction. At the population level, delay activity can differentiate between correct and error trials – a pattern that is consistent with action preparation and planning. In addition, a portion of neurons with direction selectivity can encode taste and taste recategorization. Together, these findings confirm the existence of task-related activity during the delay period and suggest that GC multiplexes information related to taste recategorization and action planning.

### Involvement of GC in the performance of a taste-based 2-AC task

Recent experimental evidence highlights that neural activity recorded in multiple brain regions, including sensory and motor cortices, correlates with movement and goal-directed behavior [22–24]. However, not all areas are instrumental for performing the task [22]. To evaluate whether the modulation of activity described above is necessary to optimally perform a taste-based 2-AC task, we silenced the GC using two experimental strategies. First, we adopted a chemogenetic approach. Adeno-associated viral (AAV) constructs (AAV8-hSyn-hM4Di-mCherry) carrying the inhibitory Gi-DREADD (hM4Di) were bilaterally injected into GC (**Figure 4A**, **Supplementary Figure 1B**). Neurons expressing hM4Di can be silenced by clozapine N-oxide (CNO) [25]. In our experimental conditions, intraperitoneal injection of CNO (10 mg/kg), but not saline (0.9%; control) significantly impaired behavioral performance (fraction of correct trials, saline vs CNO: 0.82 ± 0.02 vs 0.69 ± 0.03, paired t-test, t_(5)_ = 3.26, p = 0.02, **Figure 4B** left panel). In contrast, CNO did not affect the performance in a separate group of mice that received an injection of a control viral construct (AAV8-hSyn-mCherry) lacking the inhibitory Gi-DREADD (CNO vs saline; 0.80 ± 0.02 vs 0.79 ± 0.02, paired t-test, t_(4)_ = 0.36, p = 0.74, **Figure 4B** right panel). These results indicate that GC activity is required to perform the taste-based 2-AC task.

**Figure 4.**
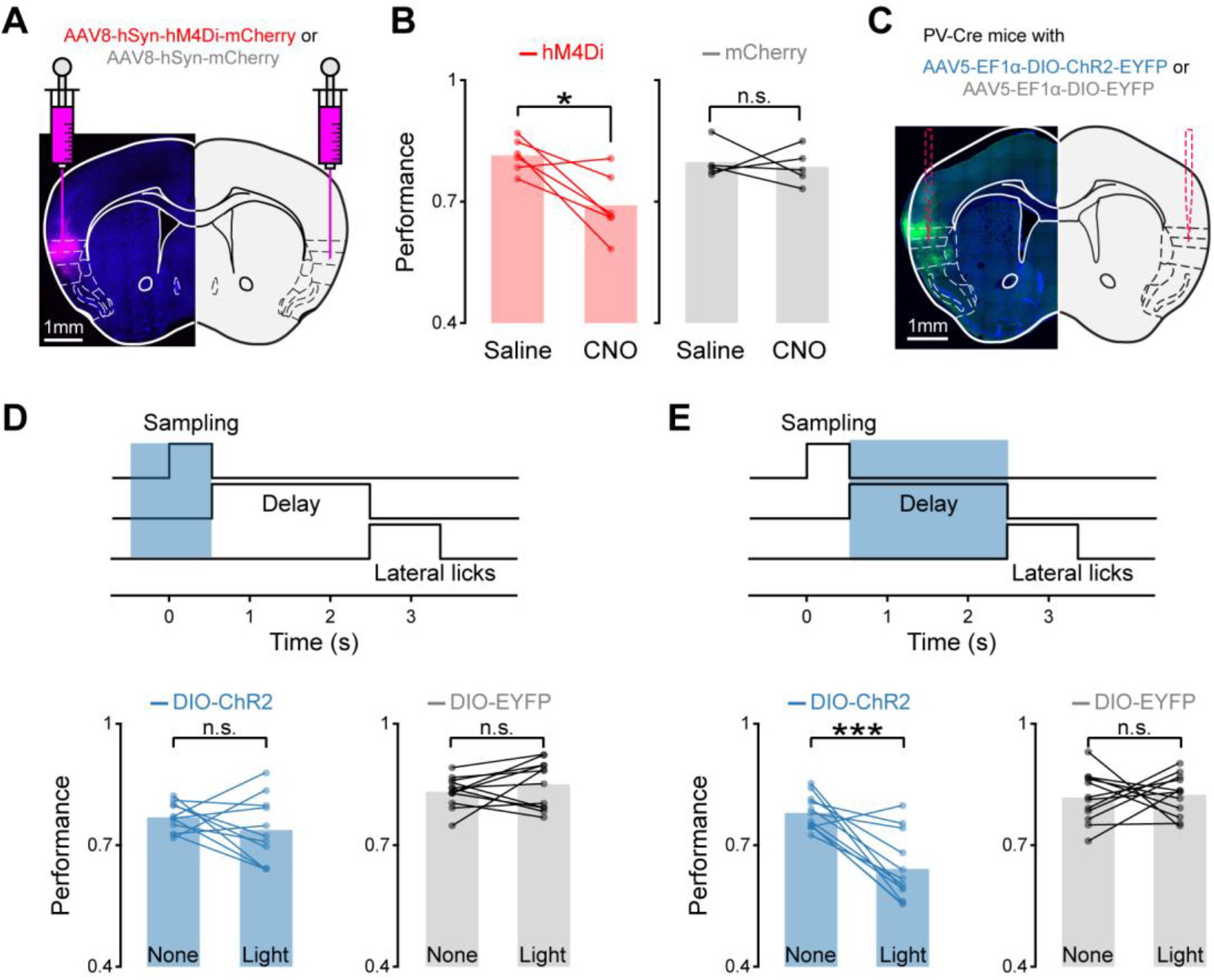
Behavioral effects of GC silencing. **A**, Sample histological section showing expression of hM4Di-mCherry (magenta) in GC. **B**, Behavioral performance (fraction of correct trials) after an i.p. injection of saline or CNO in mice with hM4Di-mCherry expression in GC (left, red, n = 6) and only with mCherry expression in GC (right, gray, n = 5). Bar plots: mean value of the performance. Paired t-test, * p<0.05, n.s. not significant. **C**, Sample histological section showing the expression of ChR2-EYFP (green) in GC and the track of the tapered fiber optic cannula. **D**, Top panel, schematic of trial structure and period of photostimulation (1 s, covering the sampling epoch). Bottom panel, behavioral performance without and with light stimulation in PV-Cre mice injected in GC with ChR2-EYFP (left, blue, 11 animal-session pairs), and with a control construct (EYFP; right, gray, 12 animal-session pairs). Bar plots: mean value of the performance. Paired t-test, n.s. not significant. **E**, Top panel, schematic of trial structure and period of the photostimulation (2 s long, covering the delay epoch). Bottom panel, behavioral performance in experimental (left, blue, 12 animal-session pairs) and control PV-Cre mice (right, gray, 12 animal-session pairs). Bar plots represent the mean value of the performance. Paired t-test, *** p<0.001

GC could be involved in mediating the performance of a 2-AC task for either its role in representing taste identity – a process predominantly happening during the sampling epoch – or for its ability to process task-related variables such as recategorization of tastants and action planning – both occurring during the delay epoch. To investigate this, we employed an optogenetic approach to transiently inhibit the GC during different epochs. AAV constructs (AAV5-EF1α-DIO-ChR2-EYFP) carrying Cre-dependent channelrhodopsin-2 (DIO-ChR2) were injected bilaterally into the GC of PV-Cre mice, resulting in the expression of ChR2 in parvalbumin (PV) expressing inhibitory neurons (**Figure 4C**, **Supplementary Figure 1C**). Optical stimulation of PV neurons is widely used to inhibit cortical circuits [18, 26–28]. Bilateral photoinhibition of GC over the sampling epoch did not significantly affect task performance (no stimulation [none] vs light stimulation [light], 0.77 ± 0.01 vs 0.74 ± 0.02, paired t-test, t_(10)_ = 1.12, p = 0.29, **Figure 4D**), nor sampling duration and reaction time (**Supplementary Figure 4B-C**). In contrast, bilateral photoinhibition of GC during the delay epoch significantly reduced the performance (no stimulation [none] vs light stimulation [light], 0.78 ± 0.01 vs 0.64 ± 0.02, t_(11)_ = 5.10, p < 0.001, **Figure 4E**) and slightly increased reaction time (1.96 ± 0.02 vs 2.03 ± 0.02, t_(11)_ = 2.94, p = 0.01, **Supplementary Figure 4F**). In a second group of PV-Cre mice where only EYFP was expressed in GC PV neurons, there was no change in performance following light stimulation during either the sampling or the delay epoch (sampling epoch: 0.82 ± 0.02 vs 0.83 ± 0.01, paired t-test, t_(11)_ = −0.23, p = 0.82; delay epoch: 0.83 ± 0.01 vs 0.85 ± 0.02, paired t-test, t_(11)_ = −1.03, p = 0.33, **Figure 4D-E**).

Altogether, these results demonstrate that GC is required for properly performing a taste-based 2-AC task, and that task performance is affected by silencing activity in the delay period, but not in the sampling epoch.

## DISCUSSION

The results presented here demonstrate the involvement of GC in a taste-guided, reward-directed decision making task. We trained mice in a taste-based 2-AC. Subjects had to sample from a central spout one out of four tastants (S, Q, M, SO) randomly selected at each trial, wait during a delay period and respond by licking one of two lateral spouts. Mice were trained to lick left in response to S and Q, or right in response to M and SO; correct responses were rewarded with water. The separation of sampling, delay and response in distinct epochs allowed us to study the temporal evolution of neural activity and its relationship to the task. We found that GC neurons represent gustatory information and task-related variables. Taste processing was not limited to the sampling epoch, but continued throughout the delay period, shifting from representing the chemical identity of tastants to representing their predictive value (lick left or right). This change in similarity of responses to S, Q, M and SO is consistent with the notion that GC dynamically recategorizes tastants according to the action they predict. Analysis of activity during the delay epoch showed that in addition to processing taste, GC neurons fired in anticipation of a licking direction, with some neurons selectively anticipating either left or right licks. Decoding analysis of correct and error trials revealed that activity in GC was not just linked to taste recategorization, but also to action preparation and planning. Indeed, responses to the same tastants differentiated correct from error trials during the delay epoch. Altogether, these recordings show that while activity in the sampling period is mostly linked to chemosensory processing, activity in the delay period reflects a recategorization of gustatory cues and preparation for a specific behavioral response. To test for the behavioral role of GC and its neural activity during the different epochs, we relied on chemogenetic and optogenetic manipulations. Silencing of GC with inhibitory DREADD led to a reduction in the overall performance, with fewer correct responses. Temporally restricted silencing of GC (by optogenetic activation of GABAergic neurons) demonstrated that silencing during the delay period significantly reduced task performance, while interfering with activity during the sampling epoch had no visible impact on behavior. Taken together, we demonstrated that the contribution of GC in a decision making task is largely due to the integration of perceptual and cognitive signals rather than just sensory processing. This result goes against classic views of cortical taste processing and emphasizes the role of GC in driving behavior.

### Temporal dynamics in GC

A well-established model of taste processing posits that GC represents taste through time varying modulations in spiking activity. In its original instantiation, this model describes the evolution of taste responses through three distinct temporal epochs unfolding over a few seconds from the delivery of a tastant [5]. The first epoch (somatosensory) lasts a few hundred milliseconds after stimulus onset and corresponds to the general tactile sensation of tastants contacting the tongue. The second epoch (chemosensory) starts after the first, lasts about a second and corresponds to a phase in which taste qualities are maximally differentiated. The third epoch (palatability) begins about a second after stimulus delivery and relates to the processing of taste palatability. This coding scheme has been further refined through trial-by-trial ensemble analyses and has been extensively validated by experimental evidence in rats and mice [6, 7, 16, 29, 30]. Alas, one of the limitations of this model has been its exclusive reliance on experiments in which rodents consume tastants that are flushed directly into the oral cavity through a surgically implanted intraoral cannula. Our experiments reaffirm and significantly expand this body of work, demonstrating that temporal multiplexing can be observed also in the context of mice engaged in a decision making task that relies on licking. We observed that chemosensation gave way to recategorization and action planning as activity progressed from the sampling through the delay epoch. Taste recategorization consisted in shifting the pairwise representation of tastants toward similarities in predicted actions (lick left *vs* lick right). Planning related signals consisted in activity which was predictive of the same licking direction regardless of the gustatory cue. Recategorization and planning were not isolated in different temporal windows, but rather intertwined during the delay epoch, suggesting that perceptual and decisional processes do not segregate in time. It is worth noting that this dynamic processing was not achieved through the activation of mutually exclusive neurons, as the same units could process multiple sensory and task-related variables (**Supplementary Figure 2B**). This result argues against the existence of cognitive labeled lines in GC.

In summary, our results demonstrate that, while the specific temporal structure and the variables encoded in GC firing rates may vary from task to task and depending on experimental conditions, the temporal multiplexing of sensory and cognitive signals is a fundamental mode of function of GC.

### Functional role of GC

GC has been implicated in multiple functions related to taste processing, taste learning and taste expectation [1, 9, 31, 32]. Recent evidence also suggests that GC can be involved in taste-based decision making [16, 33]. Recordings from GC of rats consuming tastants delivered through an intraoral cannula demonstrate that sudden and coherent changes in ensemble activity predict gapes – an innate orofacial behavior aimed at expelling aversive tastants [16]. Optogenetic experiments, showing that silencing GC prior to this transition in activity delays the onset of gapes, confirm the importance of this area in driving this ingestive decision. While important and novel, the work described above has focused exclusively on innate, ingestive responses evoked by aversive stimuli. A recent set of electrophysiological experiments relied on a 2-AC task to investigate GC activity related to decision making in the context of a structured, reward-oriented paradigm [33]. While GC showed patterns of activity consistent with decision making, it appeared less engaged by the task than the orbitofrontal cortex, raising the possibility that task-related activity might be epiphenomenal in GC. Evidence in the rodent’s brain of global preparatory signals [22] that are not necessarily instructive of behavior further raises questions on the role of reward-related, decision making activity in GC. Our experiments were explicitly designed for an in-depth investigation of patterns of firing activity associated with a 2-AC, and for a test of their behavioral significance. The reliance on restrained subjects and the use of a delay period before the decision allowed us to record task-related signals in the absence of overt movements associated with a 2-AC in freely moving rodents. Manipulation of GC activity unveiled a role for GC activity in the 2-AC task. Chemogenetic silencing resulted in a significant reduction of performance, pointing at GC playing a role in the execution of the task. Temporally restricted optogenetic silencing (through activation of PV-positive GABAergic neurons) allowed us to investigate the contribution of GC activity in different epochs, parsing apart the role of sensory and task-related signals. Silencing GC around the sampling epoch – a time in which chemosensory processing occurs with little or no cognitive signaling - had no impact on behavioral performance. On the contrary, silencing during the delay epoch – a window during which we observed firing related to taste recategorization and licking direction planning – significantly reduced the performance.

In summary, the ineffectiveness of silencing during the sampling epoch indicates that the contribution of GC to a taste-based, 2-AC is not in merely detecting gustatory stimuli at the time of licking. Nevertheless, our results point at the importance of the integration of perceptual (recategorization) and cognitive (planning) activity during the delay epoch for reward-related licking decisions. Altogether the data presented here demonstrate that the function of GC goes beyond chemosensory processing and beyond controlling the timing of naturalistic, aversive reactions, as it is also engaged in reward-related decision making.

## ACKNOWLEDGMENTS

The authors would like to acknowledge Dr. Arianna Maffei, Dr. Daniel B. Polley, Dr. Craig Evinger, Dr. Joshua L. Plotkin and the members of the Fontanini, Maffei and Vincis laboratories for their feedback and insightful comments. This work has been supported by National Institute on Deafness and Other Communication Disorders Grants R21-DC016714 to RV, R01-DC015234 and R01-DC018227 to AF.

## AUTHOR CONTRIBUTION

R.V., K.C. and A.F. carried out study conceptualization and experimental design. R.V. and K.C. performed electrophysiological recordings, behavioral experiments and data analysis. L.C. and J.C. performed behavioral experiments. All the authors contributed to writing the manuscript.

## MATERIALS AND METHODS

### Experimental subjects

Experiments were performed on 24 adult male mice (10-20 weeks old). Only male mice were used to limit the potential variability that may be introduced by estrous cycle in female mice. Sixteen C57BL/6 mice (Charles River) were used for electrophysiological recordings and chemogenetic experiments. Eight PV-Cre mice (The Jackson Laboratory, Stock # 017320) were used for optogenetic experiments. Mice were group housed and maintained on a 12 h light/dark cycle with *ad libitum* access to food and water unless otherwise specified. All experimental protocols were approved by the Institutional Animal Care and Use Committee at Stony Brook University, and complied with university, state, and federal regulations on the care and use of laboratory animals.

### Adeno-associated viral constructs

For chemogenetic experiments, we used the following viral constructs: AAV8-hSyn-hM4Di-mCherry (7.4 × 10^12^ vg/ml, UNC vector core or Duke Viral Vector Core) and AAV8-hSyn-mCherry (2 × 10^13^ vg/ml, Duke Viral Vector Core). For optogenetic experiments, we used AAV5-EF1α-double floxed-hChR2(H134R)-EYFP-WPRE-HGHpA (7.7 × 10^12^ vg/ml, Addgene, catalog #: 20298-AAV5) and AAV5-EF1α-DIO-EYFP (1.3 × 10^13^ vg/ml, Addgene, catalog #: 27056-AAV5).

### Surgical procedures for viral injections, fiber optic cannulae and electrodes implantation

Mice were anesthetized with an intraperitoneal injection of a cocktail of ketamine (70 mg/kg) and dexmedetomidine (1 mg/kg). Once fully anesthetized, they were placed on a stereotaxic apparatus. The depth of anesthesia was monitored regularly via visual inspection of breathing rate, whisking and by periodically assessing the tail reflex. A heating pad (DC temperature control system, FHC, Bowdoin, ME) was used to maintain body temperature at 35°C. Once a surgical plane of anesthesia was achieved, the animal’s head was shaved, cleaned and disinfected (with iodine solution and 70% alcohol) and fixed on a stereotaxic holder. For viral injections, a small craniotomy was bilaterally drilled above GC (AP: +1.2 mm, ML: ±3.5 mm relative to bregma). A pulled glass pipette front-loaded with the viral constructs was lowered into GC (−2.0 mm from brain surface). 100-150 nl of virus was injected at 1 nl/s with a microinjection syringe pump (UMP3T-1, World Precision Instruments, Sarasota, FL). Following injection, we waited additional 5 minutes before slowly pulling the pipette out. For optogenetic experiment, two tapered fiber optic cannulae [34] (Ø 200 μm core, emitting length = 1 mm, NA = 0.39, Optogenix, Lecce, Italy) were slowly lowered into GC (−1.85 mm from the brain surface) after virus injections (**Supplementary Figure 1C**). For electrophysiological experiments, craniotomies were opened above the left GC (AP: 1.2 mm, ML: 3.5 mm relative to bregma) and above the visual cortex for implanting movable bundles of 8 tetrodes (Sandvik-Kanthal, PX000004) and ground wires (A-M system, Cat. No. 781000), respectively. During surgery, tetrodes and reference wires (200 kΩ - 300 kΩ for tetrodes and 20 kΩ - 30 kΩ for reference wires) were lowered above GC (1.2 mm below the cortical surface). Movable bundles were further lowered 300 μm before the first day of recordings and ~80 μm after each recording session. Tetrodes, ground wires and a head screw (for the purpose of head restraint) were cemented to the skull with dental acrylic (Hygenic Perm Reline, Coltene). Before implantation, tetrodes were coated with a fluorescent dye (DiI; Sigma-Aldrich), which allowed us to verify placement at the end of each experiment (**Supplementary Figure 1A**). Animals were allowed to recover for a minimum of 7 days before water restriction regimen and training began.

### Taste-based, two-alternative choice task

Once recovered from surgery, mice were water restricted with 1.5 ml water daily for 1 week before training. Mice were head-restrained and trained in a custom-built setup to perform the taste-based 2-AC task, which was inspired by the object location discrimination task [18, 35]. The behavioral setup consisted of one central spout and two lateral spouts. Starting and ending position of the spouts and their speed were controlled by Zaber motors (X-LSM, Zaber) via LabView software. In addition, a movable aspiration line was used to clean the central spout by aspiring residues of the tastant drop after each trial. The central spout consisted of 5 independent metal tubes, each one connected to its taste line. Gustatory stimuli (sucrose [100 mM], maltose [300 mM], quinine [0.5 mM] and sucrose octaacetate [0.5 mM], Sigma-Aldrich) were delivered in ~2 μl droplets by a gravity-based taste delivery system. The lateral spouts consist of two metal tubes and were used to deliver a drop of water (~3 μl) as reward. The tips of two lateral spouts were spaced 5 mm apart from each other. Licking signals were detected with licking detectors [36], which were activated by the tongue’s contact with the metal spouts.

Mice were trained to associate sucrose (S) and quinine (Q) delivered from the central spout with water reward at the left lateral spout, and to associate maltose (M) and sucrose octaacetate (SO) delivered from the central spout with water reward at the right lateral spout. At each trial, the central spout containing a preformed drop of a tastant (pseudo-randomly chosen from S, M, Q and SO) moved close to the mouse, and started to retract once licking to the central spout was detected. This configuration resulted in a short window for sampling (~500 ms). After a delay period (average interval between the last lick for the center spout and the first lick for a lateral spout was 2 s), two lateral spouts advanced, allowing the mouse to make a lateral lick and report the choice. The first lick to either of the lateral spouts was counted as the choice. A correct lateral spout choice triggered a drop of water, while an incorrect choice triggered a time out (5 s) before the onset of the inter-trial interval. A timeout before the inter-trial interval was also triggered if the mouse failed to sample the tastants from the central spout, or failed to lick to either one of the two lateral spouts. The inter-trial interval was 6 ± 1 s.

To minimize the influence of non-gustatory cues (valve clicks, odor of tastants) on animal’s performance, experimental precautions were adopted. A fan was used to blow away the possible odor of tastants, and constant white noise was played to mask the sound of valve clicks. In addition, control experiments were performed to verify the reliance on gustatory cues in the performance of the task. A group of well-trained mice (>75% correct choices for more than 3 days in a row; n= 5) was tested in a behavioral session in which gustatory stimuli were replaced with water. Under these conditions, performance dropped to chance level (water vs tastants, 0.530 ± 0.035 **vs** 0.862 ± 0.031, paired t-test, t_(4)_ = −6.15, p = 0.003), confirming that taste information was essential to discriminate the four gustatory stimuli.

### Electrophysiological recordings

Single units were recorded via a multichannel acquisition processor (MAP data acquisition system, Plexon, Dallas, TX) in mice performing the taste-based 2-AC task. Signals were amplified, bandpass filtered (300–8000 Hz), and digitized at 40k Hz. Single units were isolated by threshold detection, and were further sorted offline through principal component analysis using Offline Sorter (Plexon, Dallas, TX). Tetrodes were lowered ~80 μm after each recording session to avoid sampling the same neurons. In total, we recorded 214 neurons from 5 mice in 21 sessions; the average yield was 42.5 neurons per mouse and 10.2 neurons per session.

### Data Analysis

Data analysis was performed using Neuroexplorer (Plexon, Dallas, TX) and custom scripts written in MATLAB (MathWorks, Natick, MA).

### Behavioral analysis

Task performance was measured as the fraction of correct trials over the total number of correct and error trials. Error trials were defined as trials in which mice licked to the wrong lateral spout. Trials with no licking to the central or lateral spouts were excluded from analysis. Normally these trials occurred at the end of the session.

### Taste-evoked response

Single unit spike timestamps were aligned to the first lick at the central spout. Perievent rasters of individual units were used to construct perstimulus time histograms (PSTH, 100 ms bin size). Taste-selective activity was assessed by examining firing rates averaged across trials and over a 500 ms window after the first central lick. Firing rates in S, M, Q and SO trials were compared using a Kruskal-Wallis test (a neuron was deemed taste selective if p < 0.05). Only neurons showing taste selectivity were further analyzed to assess the modulation evoked by a specific tastant. For each tastant, mean firing rates in a 500 ms window after the first lick to the central spout were compared with mean firing rates in a 500 ms window prior to the first lick to the central spout using a Wilcoxon rank sum test (a neuron was deemed responsive to a certain tastant if the p < 0.01).

### Population decoding of taste information

To characterize the temporal dynamics of gustatory processing in GC, we first applied a population decoder (Neural Decoding Toolbox, www.readout.info) [37]. Neurons recorded across different sessions were used to construct a pseudo population. The results presented are from 181 out of 214 neurons, as only neurons with at least 30 trials for each tastant were used to ensure robustness of classification. The results were confirmed when we relaxed the trial number constraint to 11 and included all neurons (n = 214). Spike timestamps for each neuron were aligned to the first lick of the central spout (time 0) and were binned (bin size = 100 ms) to construct a firing rate matrix, where each row represents a trial and each column represents a bin. The matrix is composed of spikes occurring from time 0 to time 2.5 s. Firing rates were normalized to Z-scores. Data were randomly divided into 10 splits, out of which 9 were used to train the classifier (max correlation coefficient) and the remaining 1 was used to test the classifier. This process was repeated 10 times, each time with different training and testing splits, to compute the decoding accuracy. Decoding accuracy within the 0-0.5 s temporal windows was averaged to represent the decoding accuracy for the sampling epoch. Decoding accuracy within the 0.5-1.5 s and 1.5 – 2.5 s temporal windows were averaged to represent the decoding accuracy during the delay. The decoding procedure was further repeated 10 times to compute the variation of the decoding accuracy for the sampling and delay epoch. In addition to the decoding accuracy, the confusion matrices within 0-0.5 s, 0.5-1.5 s and 1.5-2.5 s temporal windows were also computed.

### Visualization of population activity with principal component analysis (PCA)

To visualize the population activity, we applied PCA. Specifically, neurons recorded across different sessions (n = 214) were used to construct a pseudo population. For each neuron, spike timestamps were aligned to the first lick of the central spout (time 0) and PSTHs were computed (bin size = 100 ms, window = 0-2.5 s). A firing rate matrix was constructed for the pseudo population, where each row represents a bin and each column represents a neuron. We used PCA to find the principal component coefficients of the matrix, and applied the coefficients to the population activity evoked by S, Q, M, and SO. Population activity was projected onto the PC space. Only the first 3 PCs were used for visualization and analysis. PCA results were confirmed also when the analysis was performed exclusively on neurons with at least 30 trials for each tastant (n = 181).

### Pairwise distance between taste-evoked activities

To calculate the pairwise distance between taste-evoked activity, we applied a receiver operator characteristic (ROC) analysis for each single unit (n = 214). Single unit spike timestamps were aligned to the first lick of the central spout and PSTHs were constructed (bin size is 100 ms) for the 4 different tastants. The area under the ROC curve (auROC) was used to compute the auROC distance in neural activity between a pair of tastants: auROC_D_tastant-pair_ = | 2 × (auROC-0.5) |, ranging from 0 to 1, where 0 represents similar firing and 1 represents different firing for the pair of tastants. Distance in neural activity evoked by tastant-pairs associated with the same actions was computed as: Distance = ½ × (auROC_D_S-Q_ + auROC_D_M-SO_); and distance in neural activity evoked by tastant-pairs with same qualities was computed as: Distance = ½ × (auROC_D_S-M_ + auROC_D_Q-SO_). The results were confirmed when we only analyzed neurons with at least 30 trials for each tastant (n = 181).

### Preparatory activity during the delay epoch

Preparatory activity was first assessed only in correct trials. Single unit spike timestamps were aligned to the first lick of the lateral spout and PSTHs were constructed (bin size is 100 ms). ROC analysis [21] was then used to compare mean firing rates between left and right correct trials in a 1 s window before the first lateral lick. Specifically, the area under the ROC curve (auROC) was used to calculate the direction preference as: direction preference = 2 × (auROC-0.5). Direction preference ranged from −1 to 1, where −1 means complete preference for left trials (higher firing rate in left trials, see Neuron #1 in **Figure 3B**), 1 means complete preference for right trials (higher firing rate in right trials, see Neuron #2 in **Figure 3B**) and 0 means no preference (similar firing rate between left and right trials). To assess the significance of direction preference, we used a permutation test where left/right correct trials were shuffled without replacement. Data were shuffled 1000 times and the pseudo preference was calculated for each iteration of the shuffling. The p value was computed by comparing the actual preference with the pseudo preference. We used a criteria p < 0.01 to determine significance. Neurons with significant direction preference during the delay were defined as preparatory neurons, and the activity during the delay was deemed as preparatory activity.

Preparatory neurons were further analyzed to extract information about taste selectivity. For assessing taste selectivity, we compared activity between S and Q trials (left trials), or activity between M and SO trials (right trials) during the delay epoch (1 s before first lateral lick). We used a similar ROC analysis to quantify taste selectivity, calculated as: taste selectivity = | 2 × (auROC-0.5) |, ranging from 0 to 1, where 0 represents no selectivity between tastants (similar firing rates between S and Q trials, or between M and Q trials) and 1 represents high selectivity between tastants. We used the same permutation procedure described above to test for significance of taste response selectivity. A neuron was deemed to be taste-selective during the delay epoch if it showed either significant selectivity between S and Q or between M and SO trials. To compare taste selectivity and direction preference for each neuron, the maximum selectivity between the two pair of tastants was used (**Figure 3F**).

### Classification of correct and error trials

To analyze the relationship between preparatory activity and actions, we applied the population decoder mentioned above to the classification of correct and error trials. Preparatory neurons recorded across sessions (49 out of 89 neurons, only neurons with at least 10 error trials for both left and right trials were used) were grouped to construct a pseudo population. Spike timestamps for each neuron were aligned to the first lick of the lateral spout (time 0) and binned (bin size = 100 ms) to construct a firing rate matrix, where each row represents a trial and each column represents a bin. The matrix was composed of spikes occurring from time −2 to time 1 s. Firing rates were normalized to Z-scores. Data were randomly divided into 10 splits, out of which 9 splits were used to train the classifier (max correlation coefficient) and the remaining 1 split was used for testing it. This process was repeated 10 times, each time with different training and testing splits, to compute classification accuracy. We first applied the decoder trained with S and Q trials (including same number of correct and error trials) to classify whether trials were correct or incorrect. We then applied the decoder trained with M and SO trials (including same number of correct and error trials) to classify the correct/error trials. The overall classification accuracy of correct/error trials was represented as the averaged classification accuracy calculated for S/Q trials and M/SO trials.

To evaluate whether classification accuracy was above chance, we first shuffled the labels for correct and error trials, then trained the decoder on shuffled data to compute the null distribution of classification accuracy. Classification accuracy with p < 0.001 was deemed significantly different from the chance (**Figure 3G**, grey bar).

In addition, we calculated the direction preference for error trials. Preparatory neurons with at least 10 error trials for both left and right trials (49 out of 89 neurons) were included in this analysis. We used the same permutation test described above to calculate the significance of direction preference in error trials. In total, 12 out of 49 (24.49%) preparatory neurons show significant direction preference in error trials (red dots in **Figure 3H**).

### Analysis of the orofacial movements

Oro-motor activity was recorded at a rate of 30 frames per second with a camera placed in front of the mouse face. Images were acquired and synchronized with recorded of neural activity by Cineplex software (Plexon, Dallas, TX) and imported in Matlab for offline analysis. Only videos of orofacial movements from sessions where neurons showed direction preference were used (16 sessions) were included in this analysis. Movements of the orofacial region for each mouse were assessed by frame-by-frame video analysis [12, 13]. Briefly, a region of interest (ROI) was drawn around the animal’s mouth. Then we computed the absolute difference of the average pixel intensity of the entire ROIs across consecutive frames around the first lateral lick (time 0, **Supplementary Figure 3**). Changes in pixel intensity values of the orofacial region were normalized to background changes in pixel intensity obtained from a second ROI drawn away from the orofacial region. This allowed us correcting for changes due to fluctuations in background light intensity. Orofacial movement was represented as change in pixel intensity. We applied the same ROC analysis described above to compute the direction preference based on the change in pixel intensity in left and right correct trials. Significance of the direction preference was inferred with the permutation test described above.

### Chemogenetic manipulation of GC

See section on “**Surgical procedures for viral injections, fiber optic cannulae and electrodes implantation**” for surgical procedures. Mice with GC neurons infected with hM4Di-mCherry (n = 6) or mCherry (n=5) were used in these experiments. After learning the task and showing stable performances (correct choices > 75%) for more than three consecutive days, mice received intraperitoneal (i.p.) injection of saline (10 ml/kg body weight) or clozapine N-oxide (CNO, 10 mg/kg, 10 ml/kg, Sigma). Drugs (saline or CNO) were administered 30-40 minutes prior to the start of the behavioral sessions. CNO was stored at −20 °C and dissolved in saline (0.9%) to reach the final concentration (1 mg/ml). CNO doses were chosen based on previously published work [38]. Behavioral performance was computed as described above and compared across days with a paired t-test.

### Optogenetic manipulation of GC

See section on “**Surgical proceduresfor viral injections, fiber optic cannulae and electrodes implantation**” for surgical procedures. PV-Cre mice with GC neurons infected with DIO-ChR2-EYFP (n = 4) or DIO-EYFP (n = 4) and with implanted tapered fiber optic cannulae were used in these experiments. A 473 nm laser (473 nm, 100 mW DPSS laser system, Opto Engine LLC) was used to deliver the light. Two 470 nm LEDs were placed in front of each mouse, delivering on/off flashes at 20 Hz. LED flashing lights acted as a background masking stimulus for the laser used for photostimulation. Only 30% of the behavioral trials were randomly stimulated with the light from the laser (20 Hz, 3~4 mW). For silencing of the sampling epoch, a 1 s long pulsing light (20 Hz) was delivered from 0.5 s before to 0.5 s after the first lick to the central spout. For silencing of the delay epoch, photostimulation was delivered for 2 s after the central lick. Each mouse received 2-3 sessions of photostimulation covering the sampling epoch, and 3 sessions of photostimulation during the delay. Sessions with stimulation covering the sampling epoch were alternated with sessions for stimulation during the delay epoch. For the various conditions (i.e., silencing during sampling, silencing during delay, experimental mice and control mice), performance was compared between trials with light on and light off using a paired t-test.

### Histological staining

Mice were deeply anesthetized with an intraperitoneal injection of ketamine/dexmedetomidine (140 mg/kg, 2 mg/kg) and were intracardially perfused with PBS followed by 4% paraformaldehyde. The brain was post-fixed with 4% paraformaldehyde overnight, cryoprotected with 30% sucrose for 3 days, and was then sectioned with a cryostat into 50 μm coronal slices. For visualizing electrode tracks or the expression of the AAV constructs, slices were counterstained with Hoechst 33342 (1:5000 dilution, H3570, ThermoFisher, Waltham, MA) using standard techniques.

