## supplementary figures for "Dynamic representation of taste-related decisions in the gustatory insular cortex of mice"

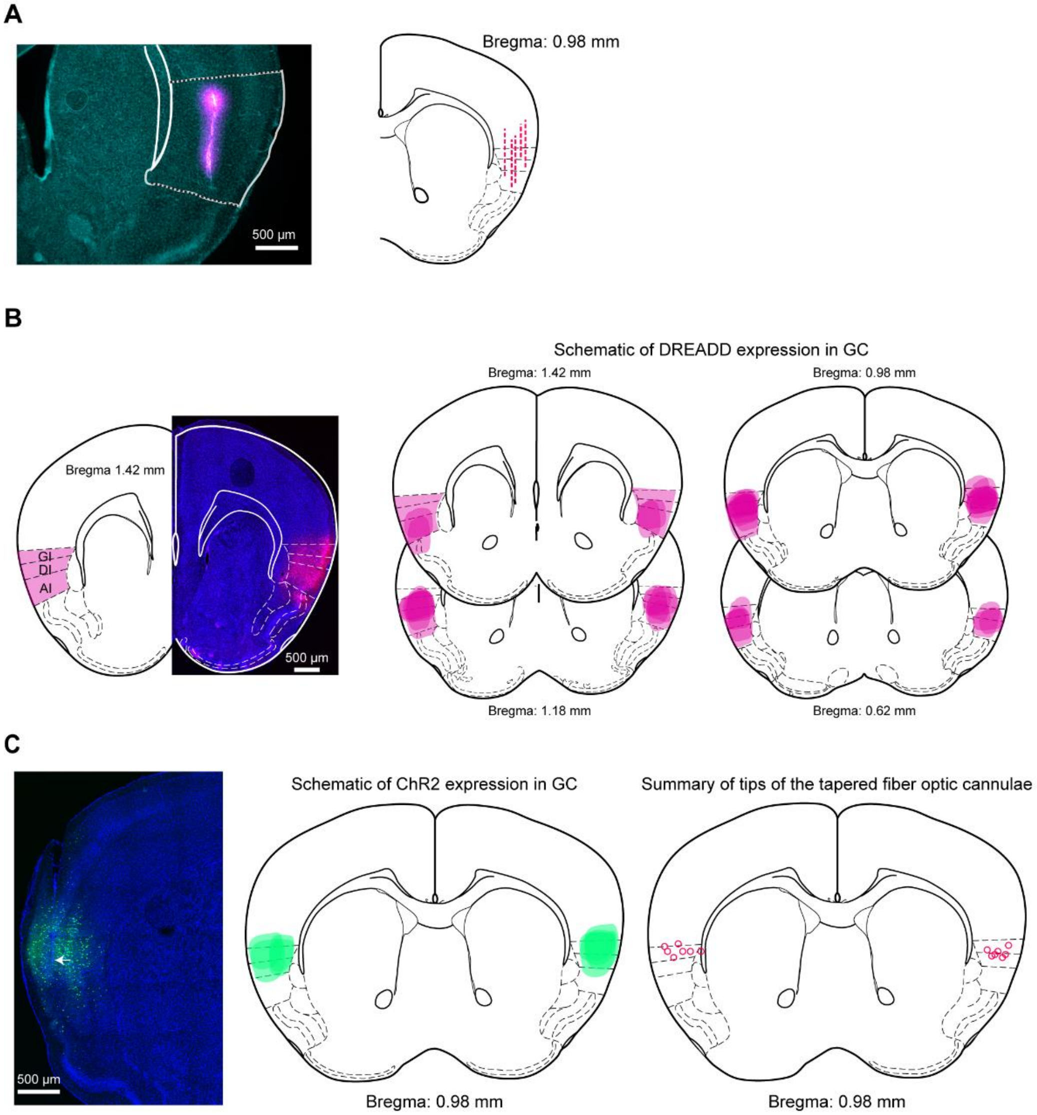


**Supplementary Figure 1. Histological verification of electrodes placement, DREADD expression, ChR2 expression and fiber optic cannulae placement. A**, Left panel: sample histological section showing a tetrode track (magenta). Right panel: schematic showing a summary of tetrode tracks. Each dashed line represents a track. **B**. Left panel: sample section showing DREADD expression (magenta) in GC. Right panel: schematic showing a summary of DREADD expression in GC from anterior to posterior. **C**. Left panel, sample histological image showing a track left by a tapered fiber optic cannula (whiter arrow), and expression of EYFP in PV neurons (green). Right panel: schematics showing a summary of ChR2 expression and tips of tapered fiber optic cannula in GC.

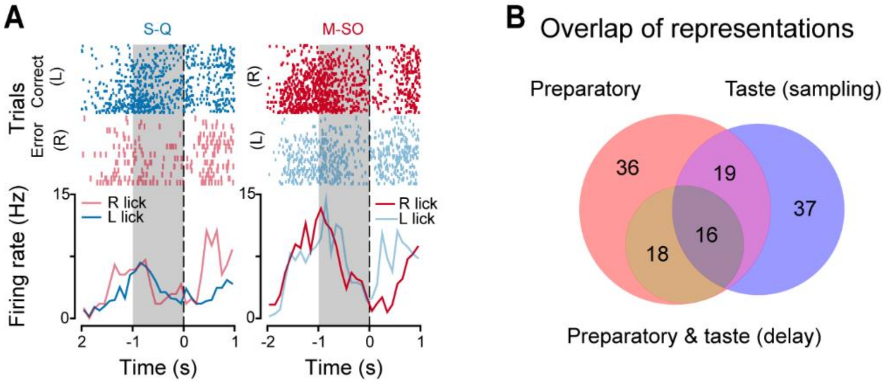


**Supplementary Figure 2. Overlap in patterns of activity. A**, Raster plots and PSTHs for a neuron showing similar neural activity during the delay epoch in correct and error trials regardless of the action. Time 0 is the first lick to the lateral spout. On the left, raster plots and PSTHs for correct (left licks, dark blue) and error (right lick, light red) trials in response to S and Q. On the right, activity for correct (right lick, dark red) and error (left licks, light blue) trials in response to M and SO. **B**. Venn diagram showing overlap of taste responses in the sampling epoch and preparatory activity during the delay epoch.

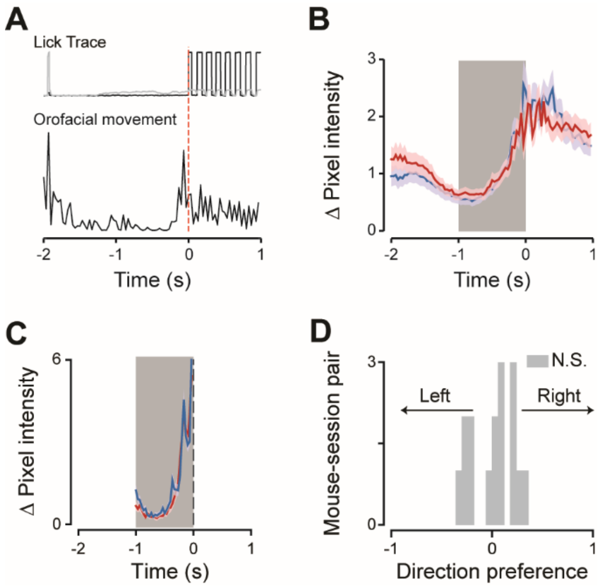


**Supplementary Figure 3. Orofacial movements and licking. A**, Representative traces of licking (grey: central lick, black: lateral licks) and orofacial movement for a single trial. Time 0 represents the first lick to the lateral spout. **B**, Time course of average orofacial movements (n = 16 sessions). Time 0 is the first lick to the lateral spout. Blue and red traces represent orofacial movements for left and right correct trials, respectively. The shaded area represents the delay period (-1 to 0 s) used for further data analysis. **C**. Time course of orofacial movements during the delay epoch (-1 to 0 s) recorded from the same sessions as the neural activity shown in **Figure 3B** Neuron #1. Blue and red traces represent orofacial movements during the delay epoch for left and right correct trials, respectively. **F**, Histogram of direction preference computed using orofacial movements data. None of the 16 sessions showed any significant direction preference (gray bars represent sessions where orofacial movement for left and right trials are not significantly different).

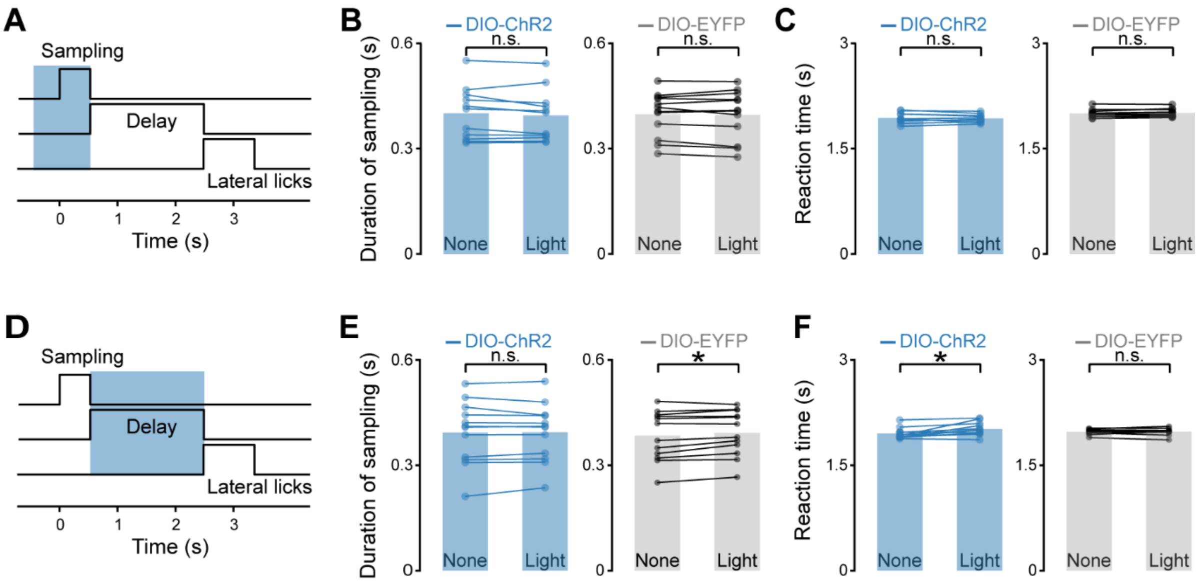


**Supplementary Figure 4. Effects of optogenetic silencing on licking. A**, Diagram of trial structure and period of photostimulation of PV neurons (1 s long, covering the sampling epoch). **B** and **C**, Duration of licking (**B**) and reaction time (**C**) without or with light stimulation during the sampling epoch in PV-Cre mice injected in GC with either ChR2-EYFP (left, blue, 11 animal-session pairs) or with a control construct (right, gray, 12 animal-session pairs). Bar plots: mean value of licking duration (**B**) and reaction time (**C**). Paired t-test, n.s. not significant. **D**, Schematic showing the trial structure and the period of the photostimulation (2 s long, covering the delay epoch). **E** and **F**, Duration of licking (**E**) and reaction time (**F**) in experimental (left, blue, 12 animal-session pairs) and control (right, gray, 12 animal-session pairs) PV-Cre mice. Bar plots represent the mean value of licking duration (**E**) and reaction time (**F**). Paired t-test, * $p<0.05$.
